# The MosAICo ecosystem: bridging the taxonomic gap in vector surveillance with real-time entomological artificial intelligence

**DOI:** 10.64898/2026.06.20.733369

**Authors:** Noemi Sarleti, Alberto Tubito, Francesco Severini, Vittorio Dante, Andrea Ciardiello, Francesco Silvestrini, Mariangela Bonizzoni, Yaw Asare Afrane, the MosAICo Working Group, Marco Di Luca, Guido Gigante, Pietro Alano

## Abstract

Mosquito-borne diseases represent an escalating global health threat, driven by climate change, urbanization, and the spread of invasive vectors into new territories. Effective surveillance is constrained by a critical ‘taxonomic impediment’: the rate of specimen collection far outpaces the capacity of expert entomologists to process and identify trap catches. To address this bottleneck we developed MosAICo, an integrated AI-powered ecosystem for automated mosquito species identification designed for real-world, national-scale entomological surveillance. The system combines a standardized benchtop imaging device with MosAICo-Net, a deep learning pipeline enabling efficient and principled open-set recognition and uncertainty quantification. Trained and evaluated on 12, 499 specimens spanning 15 species collected across Italy, the model identifies seven priority vector species while explicitly rejecting out-of-distribution specimens. On a geographically stratified held-out test set, MosAICo-Net achieved over 90% accuracy on target species, and an AUROC of 0.96 for out-of-distribution detection. Field validation across 20 Italian surveillance sites confirmed these results: 94% micro accuracy on 1, 470 field-collected target specimens and strong agreement with expert manual counts (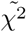 = 0.66). To assess cross-geographic generalizability, the system was further evaluated on 118 *Aedes albopictus* specimens collected at the fringe of the species invasion front in Ghana: a 97.4% accuracy with only a single specimen escalated to expert review, suggests that MosAICo is well-suited for deployment in distant and epidemiologically critical regions. The system processes up to 82 specimens per image, matching expert throughput at constant speed regardless of taxonomic complexity. By embedding uncertainty-aware AI within a standardized hardware-software pipeline, MosAICo acts as a scalable force multiplier for public health entomology, freeing expert attention for rare, invasive, or ambiguous specimens that require human validation.

## 1 Introduction

The concept of planetary health security is predicated on the stability of ecological boundaries. However, in the Anthropocene epoch, these boundaries are becoming increasingly permeable (European Centre for Disease Prevention and Control and European Food Safety Authority, 2025; Simonin, 2025). The confluence of hyper-urbanization and climate warming has fundamentally altered transmission dynamics, effectively ‘de-territorializing’ vector-borne threats. This ecological shift creates unprecedented opportunities for ancient pathogens—evolutionary lineages of viruses such as dengue, West Nile, and Zika that have coevolved with their hosts for millennia—to exploit modern infrastructure and colonize new latitudes (Hou et al., 2024; Zhang et al., 2024). The significance of mosquito-borne pathogens stems from their global ubiquity and their dual impact on human and animal health. Unlike simpler infectious models, the epidemiology of mosquito-borne diseases (MBDs) is characterized by extraordinary complexity, often requiring the navigation of intricate transmission cycles across diverse vectors and hosts. This dynamic landscape is being reshaped by a synergy of interconnected determinants: environmental shifts and climate change, alongside demographic pressures such as urbanization and the intensified global movement of people and livestock (Thomson, 2018). Within this context, land-use alterations and pathogens spill over across different hosts further complicate the emergence and re-emergence of these threats.

Addressing such challenges requires—as posited by Wilcox and Gubler (2005)—a holistic understanding of biological systems that acknowledges their inherent complexity. As invasive mosquito vectors expand into warming temperate zones, these pathogens are no longer tropical anomalies, but systemic risks within globalized urban networks. Consequently, establishing a more granular and profound comprehension of mosquito ecology and spatial distribution is imperative to refine control strategies (Wouters et al., 2024). In this shifting landscape, the management of MBDs has become a global priority, and effective mosquito surveillance and control remain the primary lever to decouple ecological change from pandemic risk.

Traditional vector surveillance strategies face considerable operational limitations, encompassing insecticide resistance and the logistical challenges of large-scale monitoring. Consequently, the conventional model of entomological surveillance—reliant on specific skills and a constrained workforce of taxonomy experts—has reached a functional limit. This has resulted in a significant ‘taxonomic impediment’, whereby the pace of environmental and ecological changes significantly outpaces the capacity for manual data processing (Høye et al., 2021; Kumar and Govindasamy, 2025; Sedda et al., 2025). Within this context, automation through systems like MosAICo becomes essential to overcome these constraints, specifically serving two critical operational goals: first, to rapidly detect the introduction of invasive mosquito species prior to their establishment; and second, to systematically map and monitor the spatial distribution of indigenous (native) and colonized (naturalized) vector species.

This paucity of experts has resulted in a triage crisis, whereby highly trained entomologists are compelled to dedicate the majority of their time to the enumeration of abundant, prevalent species (e.g., *Aedes albopictus*) rather than concentrating on rare, invasive threats or complex validations.

In order to address this bottleneck, the field of ‘Computational Entomology’ has emerged as a promising area of research, leveraging Machine Learning (ML) to automate identification across diverse modalities. Research in this field has diversified into geospatial, visual, and audio models: Convolutional Neural Networks (CNNs) have shown high efficacy in controlled settings (Abdi et al., 2025; Goodwin et al., 2021; Okayasu et al., 2019; Teixeira et al., 2023), while acoustic monitoring has been posited as a scalable alternative (Bilal et al., 2023; Mukundarajan et al., 2017; Supratak et al., 2024). Concurrently, digital surveillance has expanded through distributed IoT infrastructures, such as intelligent traps (Oliveira and Mafra, 2024) and social media mining (Melo et al., 2024).

Despite this momentum, substantial practical limitations prevent full operational integration. The challenges facing the field can be attributed to three convergent barriers. Firstly, distributed hardware networks have been shown to be challenging to scale in the face of environmental volatility (Oliveira and Mafra, 2024). Secondly, the utilization of heterogeneous digital sources frequently engenders noise rather than actionable intelligence (Melo et al., 2024). Thirdly, and most critically, approaches to computer vision face a so-called ‘robustness gap’. Most of existing models (Kasinathan et al., 2021; Goodwin et al., 2021) are designed for single-specimen identification and lack uncertainty quantification, leaving them unable to flag predictions as unreliable — whether due to poor specimen condition, suboptimal imaging geometry, or intrinsic model uncertainty — or to detect specimens belonging to species outside their training distribution (Badirli et al., 2023; Chen et al., 2025). Moreover, such systems frequently encounter difficulties when processing multiple specimens (Bjerge et al., 2023) and frequently lack the incorporation of essential quality characteristics, such as anatomical integrity, which are necessary to trust a prediction for operational decision-making. This incapacity to manage ‘open-set’ scenarios gives rise to elevated false-positive rates, thereby rendering contemporary tools unfit for secure decision-making processes.

To bridge this gap, we developed MosAICo. Rather than replacing expert entomologists, MosAICo is designed as a force multiplier: by automating the identification of routine, high-confidence specimens, it frees expert bandwidth for the ambiguous or epidemiologically critical cases that genuinely require human judgment. To achieve this objective, the ecosystem standardizes image capture in order to match the neural network’s training domain. Automated entomological surveillance requires minimizing visual noise and domain shift—such as variations in lighting, background, or camera distance—which heavily degrade neural network performance. To address this, Brey et al. (2022) introduced a foundational paradigm with the ‘MosID’ device, demonstrating that strictly controlling the physical imaging environment is a prerequisite for reliable automated identification. By standardizing illumination and hardware parameters, their approach proved that stabilizing inputs at the capture stage ensures that operational images consistently match the training domain of machine learning models. Directly adopting and building upon this very approach, the MosAICo system incorporates these identical principles of environmental stabilization to ensure consistent data input.

Crucially, MosAICo has been specifically implemented and validated in Italy, targeting the unique morphological and ecological profiles of local mosquito populations. The Italian peninsula serves as a critical case study due to the Mediterranean Basin’s status as a primary gateway for the introduction and establishment of invasive mosquito vectors Gasperi et al. (2012). This region’s geographical position, combined with its temperate climate and extensive trade networks, creates a highly favorable environment for the emergence and spread of pathogens such as chikungunya, dengue, West Nile and Usutu viruses (Mingione et al., 2023; Menegale et al., 2025; Marini et al., 2021). By deploying this system within such a high-risk ecological theater, we provide a scalable framework that not only supports national public health efforts but also serves as a robust model for vector-borne disease surveillance across the broader Mediterranean landscape. Building upon the principle of environmental standardization to address these specific ecological threats, the MosAICo ecosystem extends this control to the more complex challenge of high-throughput batch identification. Rather than limiting control to isolated or static specimens, MosAICo integrates a standardized capture pipeline with a digital-native surveillance infrastructure. By engaging academic institutions and zooprophylactic institutes—effectively replicating at the national level the network of European experts created by ECDC through VectorNet (Wint et al., 2023)—this pipeline is capable of generating standardized, real-time entomological intelligence across decentralized nodes.

## 2 Results

The foundation of the MosAICo ecosystem is a custom-built, geographically diverse image database designed to capture the true morphological variance of Mediterranean mosquito vectors. The MosAICo ecosystem integrates two tightly coupled components: a standardized hardware image acquisition unit and the MosAICo-Net inference pipeline, jointly designed to transform unstructured trap catches into structured, machine-readable entomological records (Figure 1).

**Figure 1:**
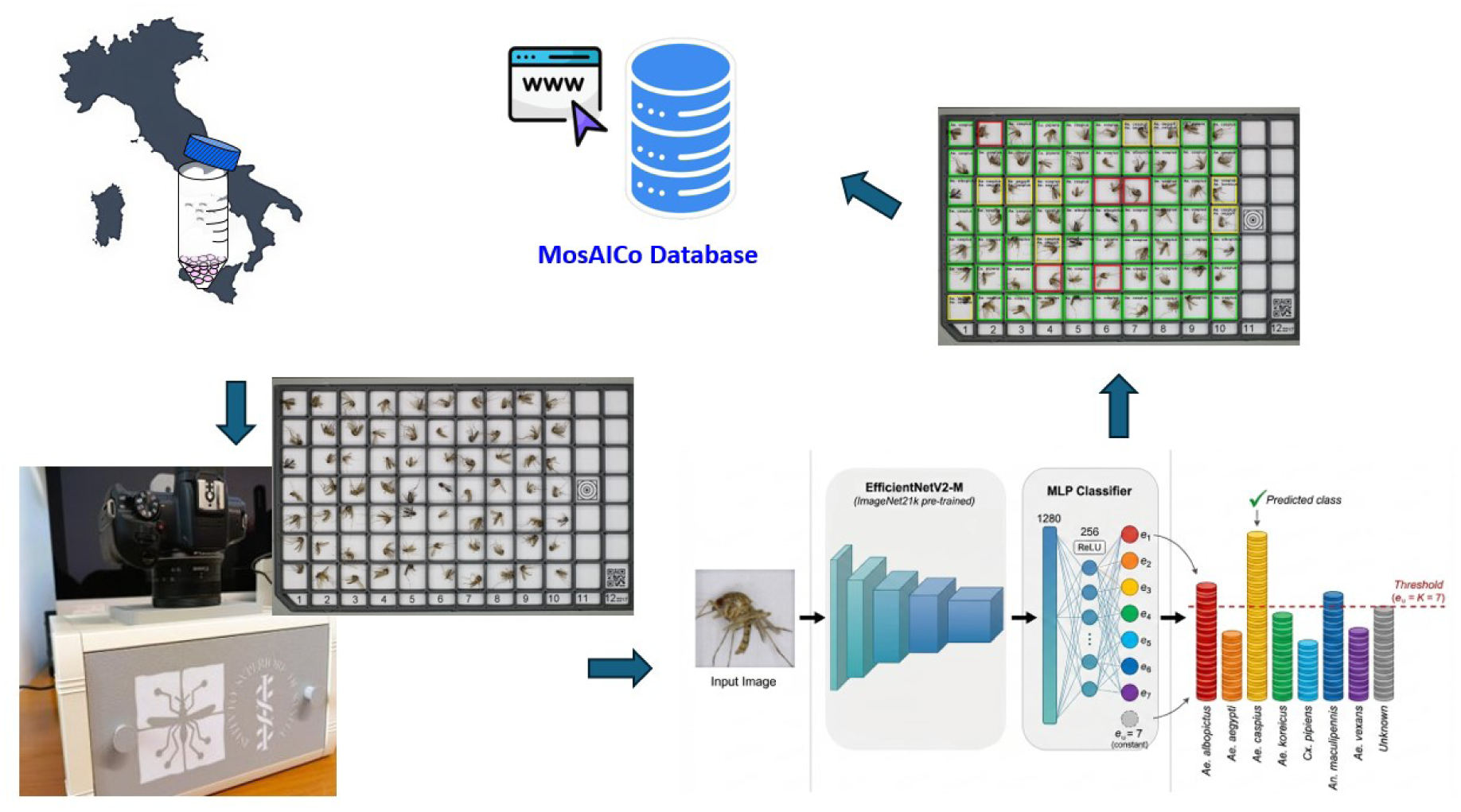
Mosaico end-to-end surveillance workflow. Mosquitoes collected in the field are placed into a grid cassette accommodating up to 82 individuals in isolated, non-overlapping compartments. The populated cassette is inserted into the MosAICo acquisition device, which automatically captures a high-resolution image and uploads it to the remote server. Server-side processing restricts analysis to occupied cells, mapping model predictions to a color-coded output: green denotes a high-confidence single-species identification; yellow flags ambiguity between two candidate species for expert review; red indicates high uncertainty, suppressing the prediction and requiring entomologist intervention. All associated metadata — such as geolocation, timestamp, and operator identity — are stored in a central database and made accessible via a web dashboard.

The algorithm was trained using specimens collected throughout Italy via the MosAICo network, sourced from both extensive field activities and established insectaries. While 25 different mosquito species were initially collected, only seven were selected for the training phase. This selection was based on high sample density and the requirement for each species to represent at least three distinct geographical origins, ensuring robust model training and diversity. The training set included *Aedes aegypti*, a species of primary concern due to its role as a potential emerging vector in Europe.

### 2.1 Deep Learning Model Performance

The model was evaluated on a geographically stratified held-out test set of 1, 577 mosquitoes, comprising both field-caught and laboratory-reared specimens. All specimens were sourced exclusively from collection sites absent from the training corpus, providing a critical assessment of the system’s generalization to previously unseen deployment environments. The two species sourced exclusively from a single collection site (*Anopheles algeriensis* and *Aedes rusticus*) were reserved for the test set, allowing for a rigorous evaluation of the model’s ability to generalize to previously unseen species.

For the seven target mosquito species, the model demonstrated high sensitivity (Figure 2), achieving 93% micro accuracy (overall accuracy across all samples) and 92% macro accuracy (average accuracy computed per species).

**Figure 2:**
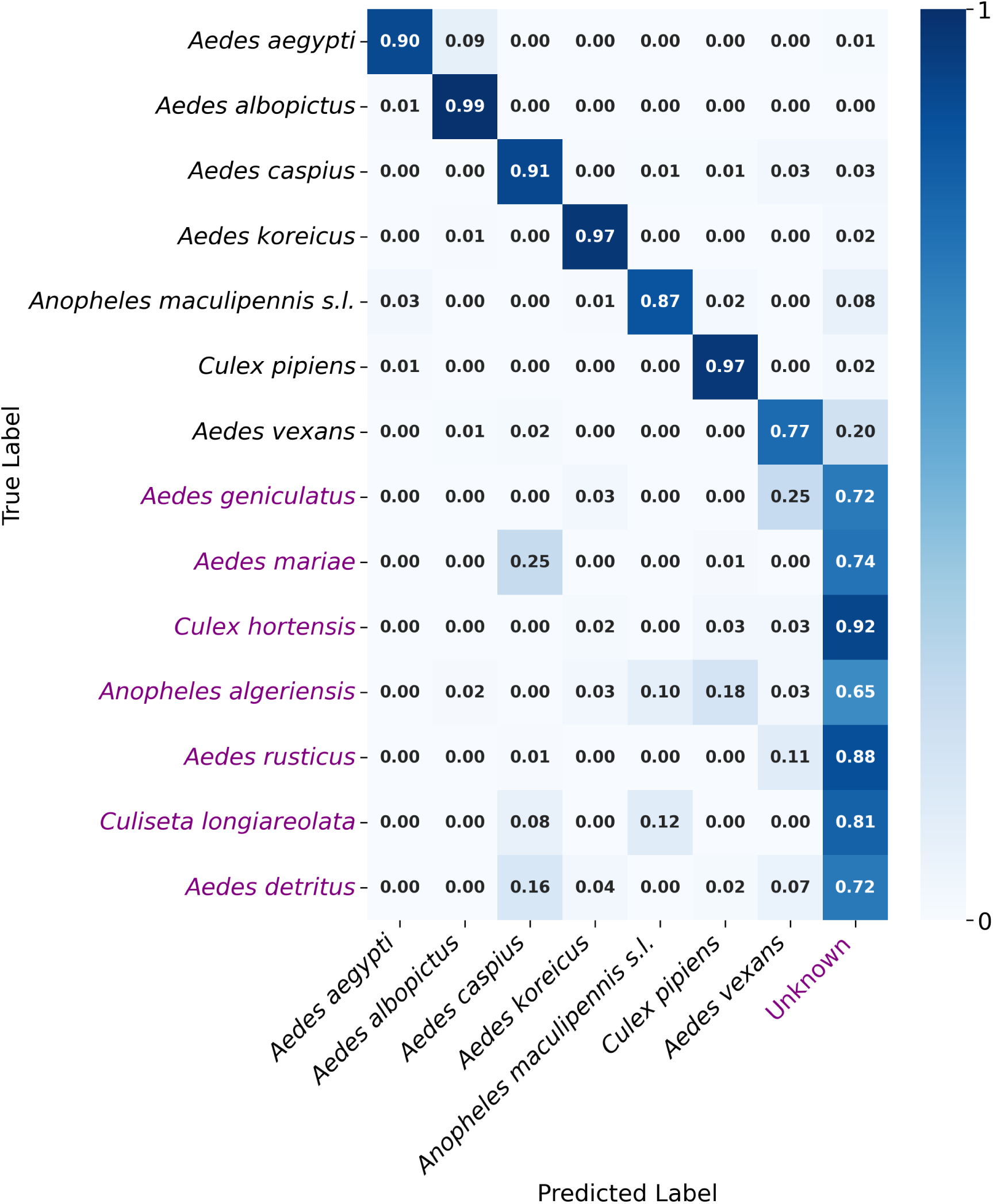
Lab test set confusion matrix. Classification performance across the 7 target species and unknown species. The matrix reveals a “block-diagonal” structure: misclassifications are heavily concentrated within genus blocks (e.g., confusing *Aedes aegypti* with other *Aedes* spp.), whereas inter-genus errors (e.g., *Aedes* vs. *Culex*) are negligible. The seven target species (bold) are: ***Aedes aegypti*** (364), ***Aedes albopictus*** (263), ***Aedes caspius*** (239), ***Aedes koreicus*** (146), ***Anopheles maculipennis sensu lato (s.l.)*** (173), ***Culex pipiens*** (284), ***Aedes vexans*** (106). Non-target species used as anchor unknowns during training are: *Aedes geniculatus* (88), *Aedes mariae* (81), *Culex hortensis* (59), *Culiseta longiareolata* (52), *Aedes detritus* (57). True out-of-distribution species, withheld entirely from training, are: *Anopheles algeriensis* (195), *Aedes rusticus* (174).

In addition to ensuring the utmost precision, the operational safety of the system was validated through its capacity to reject non-target species. By monitoring maximum target evidence, the model achieved an AUROC (Area Under the Receiver Operating Characteristic curve) of 0.96 on the test set, for Out-Of-Distribution (OOD) detection (Figure 3). It is noteworthy that high-confidence misclassifications were heavily genus-constrained (Figure 2).

**Figure 3:**
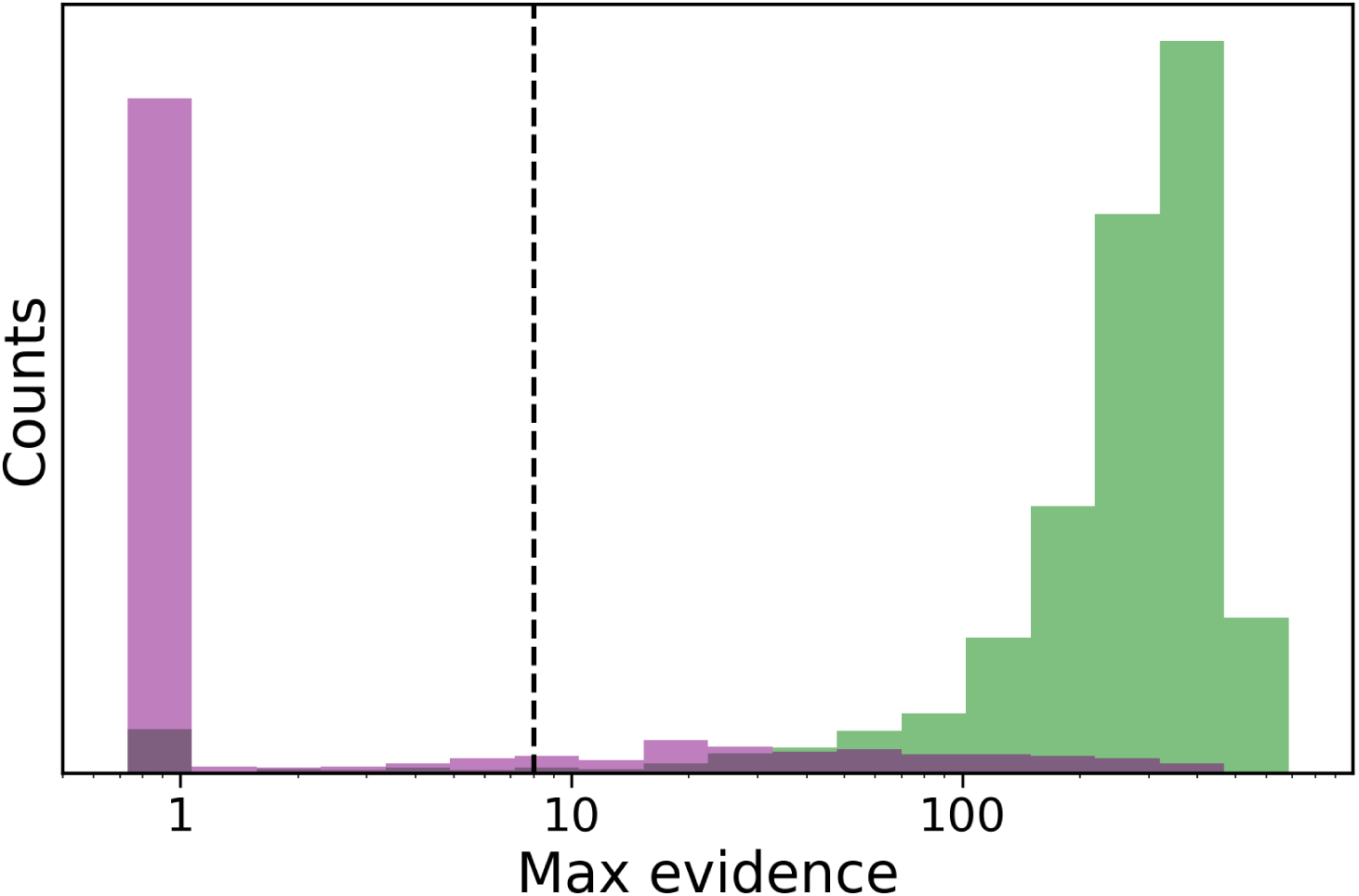
OOD detection. Distribution of top target evidence scores for Target Species (Green) vs. Non-Target (Purple). The system successfully assigns low evidence scores to non-target species, effectively filtering them as “Unknowns”. The threshold *e_u_* = 7 is represented by a black vertical dashed line.

As a macro averaged estimate, 76% of the time these errors nevertheless correctly identified the genus of the mosquitoes, while only 24% were misidentified as belonging to another genus. For example, the presence of species unknown to the algorithm led to occasional confusion with other specimens of genus *Aedes* but rarely resulted in misclassification as *Anopheles* or *Culex*. This behavior confirms a “Hierarchical Safety” mechanism. From a public health perspective, the system ensures an “operational safety” that can be defined as the capacity to function reliably while minimizing the risk of misidentification.

To interpret the learned features, we projected the high-dimensional embeddings of the training set into a 2D latent space using a Variational AutoEncoder (VAE). Visual inspection (Figure 4) revealed that unknown species formed a central cluster near the origin (0, 0), which aligns with the m-EDL interpretation that the network produces no evidence for these unknowns (i.e., the network does not activate). In contrast, each target species occupied a distinct, elongated cluster, confirming that the model relies on robust morphological features that individuate the latent direction corresponding to the variation of species evidence.

**Figure 4:**
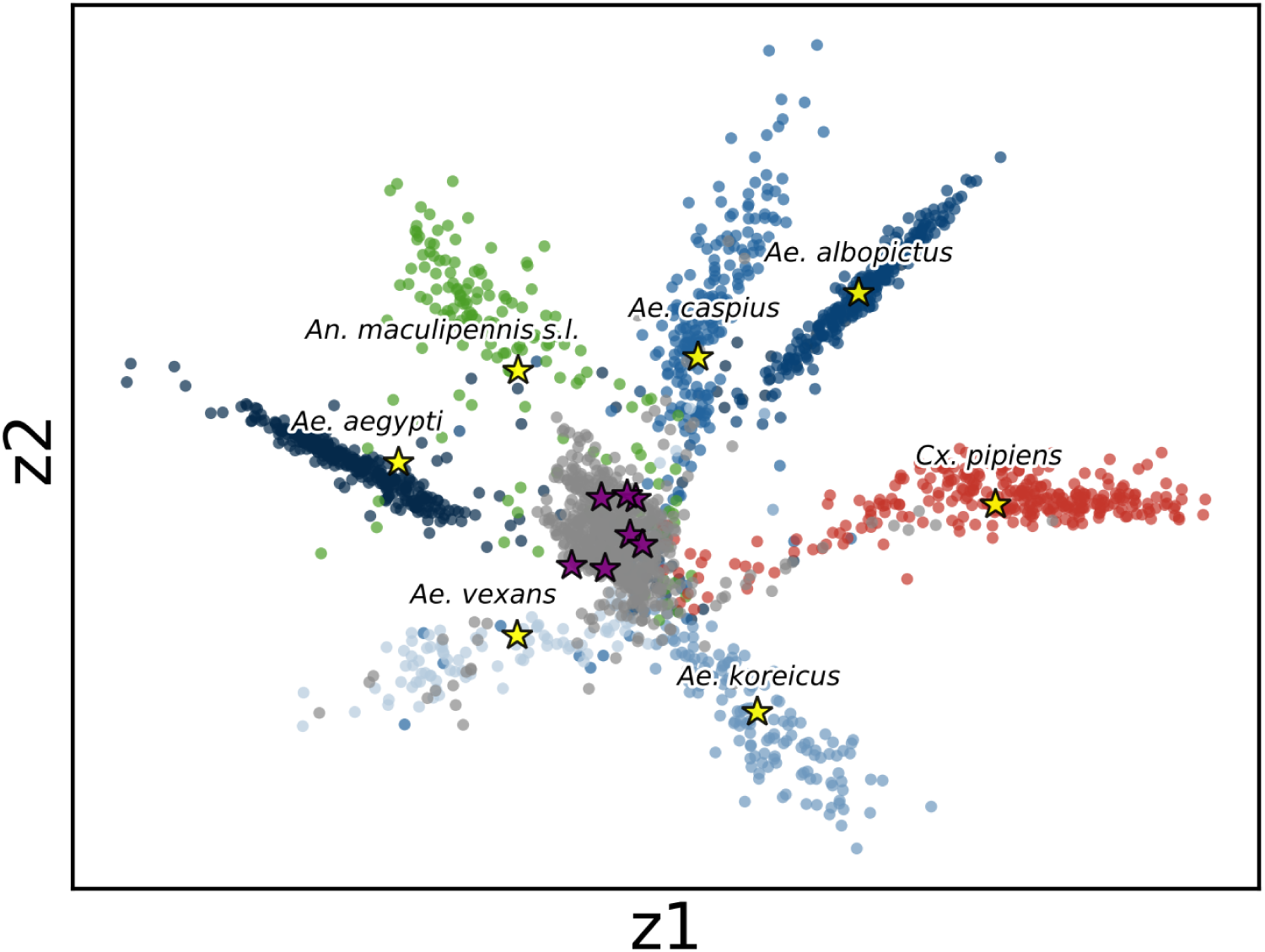
Latent space visualization. Two-dimensional projection of the feature embeddings for the test set. Target species (colored clusters) form tight, distinct islands, confirming the model learns robust morphological features. The centroids of each target species cluster are marked with yellow stars. In contrast, “Unknown” non-target species (grey points) form a scattered periphery, demonstrating their separation from the known mosquito distribution. The centroids of the different unknown species groups are denoted by purple stars.

### 2.2 Field validation of Deep Learning Model Performance

To evaluate the MosAICo system under real-world conditions, 25 high-resolution images containing 1,522 mosquito specimens were collected across Italy during the summer 2025 surveillance campaign. The deployment of ten standardized devices across 20 sites provided a radiometrically consistent baseline to rigorously test the AI-predicted counts against the gold-standard manual identifications performed by expert entomologists. The model maintained high classification accuracy, validating the decision to train on “noisy” field data. For the target species collected in the field (1470 specimens in total), the model demonstrated high sensitivity, achieving micro and macro accuracy of 94% and 89%, respectively (Figure 5). In contrast, the evaluation of performance on non-target species (52 specimens in total from 10 distinct species, 6 of which were represented by just one specimen) resulted in a micro accuracy of 35% (and a macro accuracy of 39%). This performance level constitutes only a highly approximate estimate, primarily due to the severe limitations of the field test set. Moreover, the model faced a more difficult classification challenge as only 9 of these specimens belonged to species encountered during the training phase. Nevertheless, it is important to note that, as expected, misclassifications are consistently confined to species belonging to the same genus (88% errors are genus-constrained).

**Figure 5:**
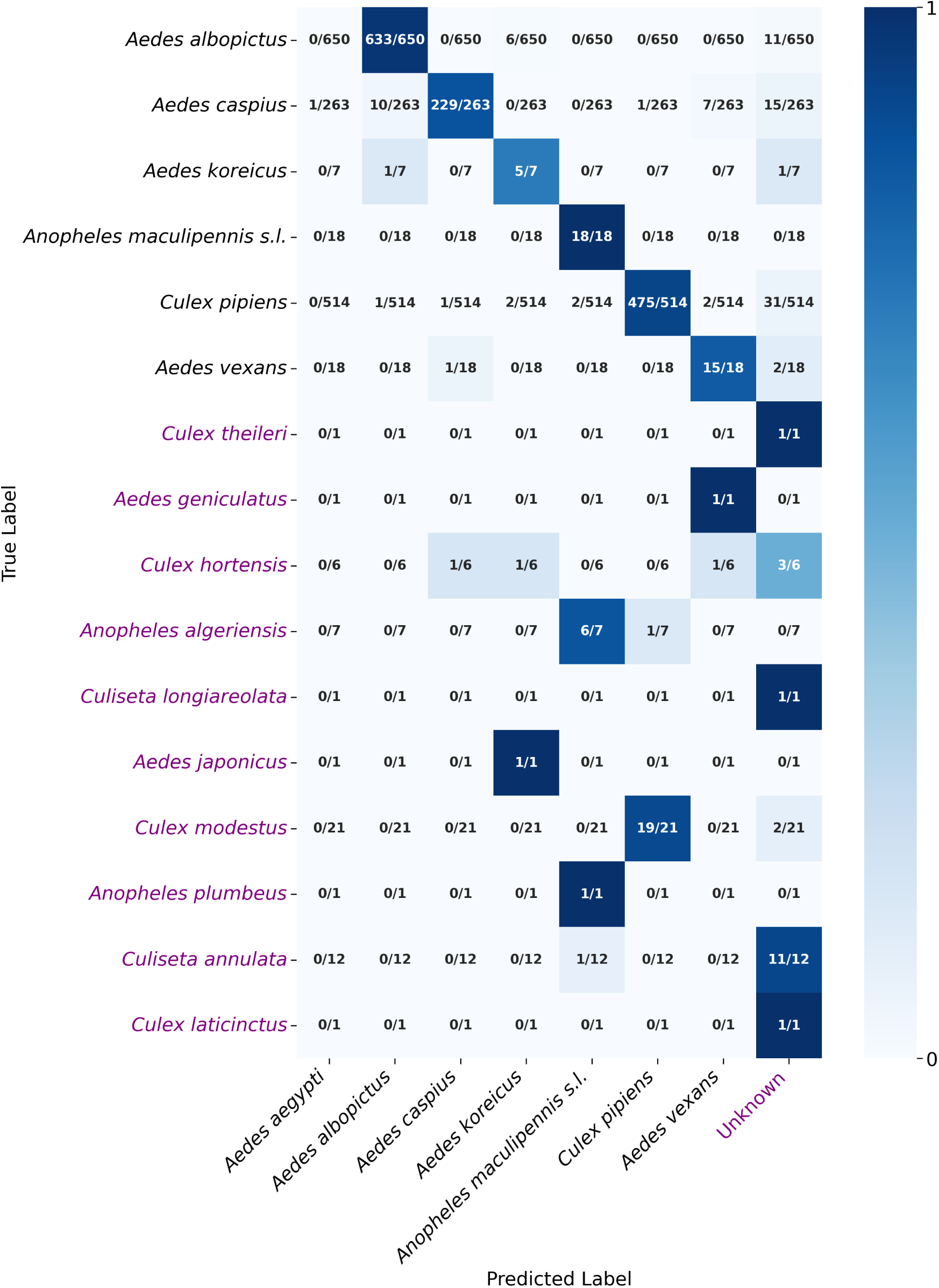
Field confusion matrix. Confusion matrix illustrating classification performance across the field-collected specimens. The field set comprised six of the seven trained target species (excluding *Ae. aegypti*, which is not currently present in Italy), and 10 non-target species, six of which were True Out-of-Distribution (OOD), never presented to the model during training. Consistent with the system’s inherent “Hierarchical Safety” mechanism, misclassifications were heavily concentrated within genus blocks (e.g., confusing *Culex pipiens* with other *Culex* spp.), while inter-genus errors (e.g., *Aedes* vs. *Culex*) were negligible.

Beyond per-specimen accuracy, population-level species composition — the quantity most directly relevant to surveillance decisions — was evaluated as the key operational metric. MosAICo is not designed to replace the entomologist, but to lighten their workload: green- and yellow-box predictions are reported autonomously, while red-box specimens are withheld and escalated for expert review, ensuring that human expertise is reserved for the small fraction of cases that genuinely require it. In field validation, such uncertain cases represented only ∼ 5% of the total catch. Agreement between automated and manual counts on the remaining 95% was assessed via a chi-squared test (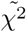 = 0.66, with a p-value of 0.68), confirming that the model accurately reproduces population-level species composition within sampling noise (Figure 6).

**Figure 6:**
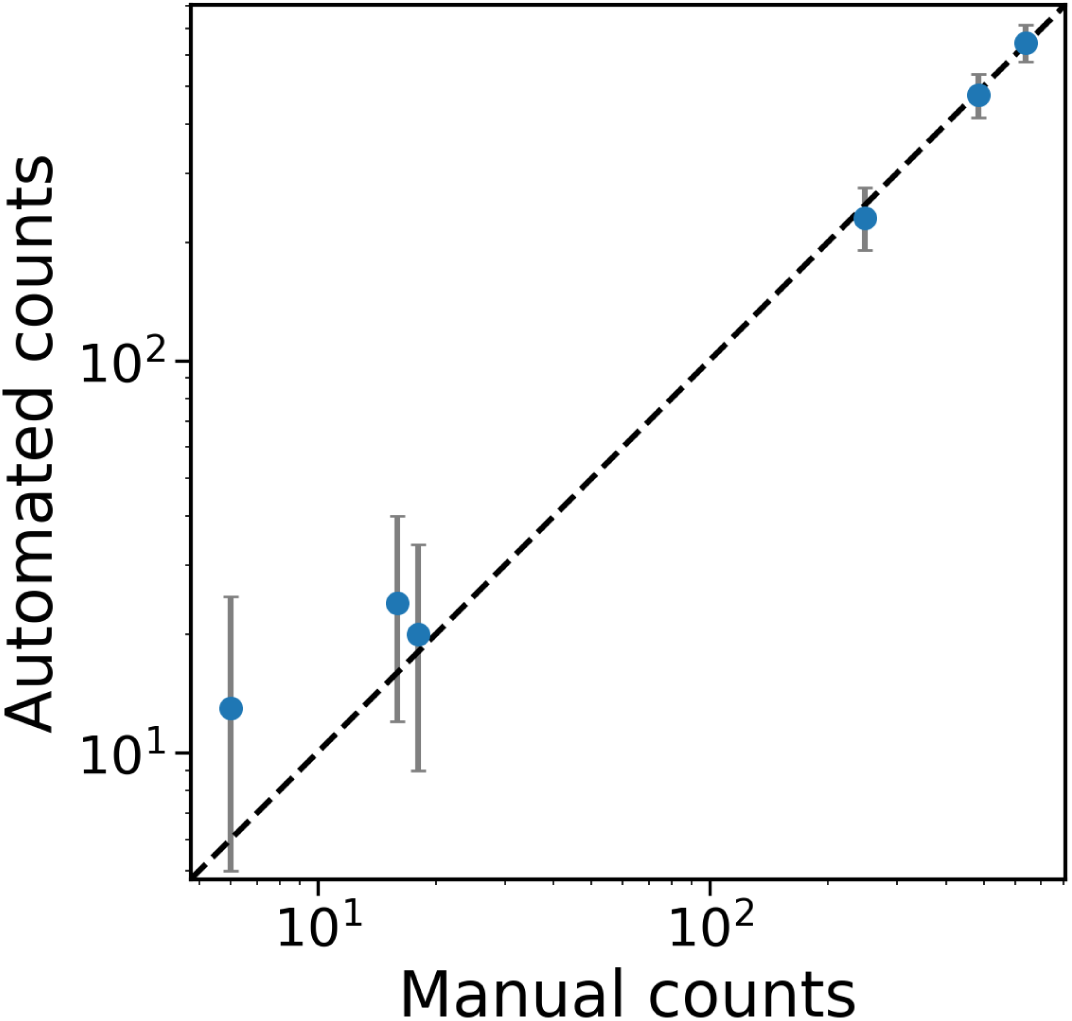
Automated counts. Comparison of MosAICo automated counts (y-axis) against expert manual counts (x-axis) for 1470 specimens belonging to six target species across the 20 validation sites. The analysis does not include *Ae. aegypti* because it is not currently present in Italy.

### 2.3 Cross-Geographic Generalization: *Aedes albopictus* from Ghana

To assess whether MosAICo-Net maintains its performance on target species from geographically distant populations outside the training distribution, the system was evaluated on 118 *Aedes albopictus* specimens from Ghana, reared under controlled insectary conditions at the University of Pavia and pre-identified by expert entomologists. No model fine-tuning or domain adaptation was performed. Specimens were organized into four imaging grids corresponding to distinct post-emergence age cohorts: 1-2 days (Grid 1, n = 35), 7-8 days (Grid 2, n = 30), 14–15 days (Grid 3, n = 30), and more than 40 days (Grid 4, n = 23).

Of the 118 specimens, 99 (83.9%) received a green box (confident single-species prediction), 18 (15.3%) a yellow box (ambiguous top-2 prediction), and only 1 (0.8%) a red box (escalated to expert). On the 99 green-box specimens, top-1 accuracy was 99.0% (98/99), with the single misclassification as *Ae. koreicus* occurring in Grid 4 (the oldest age group). Across all 117 non-red specimens, overall top-1 accuracy was 97.4% (114/117) and top-2 accuracy was 99.1% (116/117). Yellow-box ambiguities arose predominantly as *Ae. albopictus* vs. *Ae. aegypti* (12/18 cases) or *Ae. albopictus* vs. *Ae. koreicus* (6/18 cases), consistent with the intra-genus confusion pattern documented in the Italian validation.

Both the single red-box case and the elevated proportion of yellow boxes in Grid 4 suggest that aging-related morphological degradation marginally increases model uncertainty, as expected from the progressive deterioration of discriminative morphological features in older specimens.

### 2.4 Operational Throughput and Efficiency

Efficiency gains were assessed through a survey distributed to the partner network, where the MosAICo workflow was benchmarked against manual methods using a 5-point Likert scale (from 1 = ‘Strongly Disagree’ to 5 = ‘Strongly Agree’) to standardize performance feedback. Results demonstrated that the MosAICo system performs on par with an expert entomologist, processing a bulk sample of 82 specimens with an average processing time of 11.5 minutes compared to the human average time of 11.7 minutes.

Notably, while an entomologist’s processing time increases with sample heterogeneity and the presence of rare species, MosAICo maintains a consistent speed regardless of taxonomic complexity. Furthermore, the workflow was rated highly for its intuitiveness, achieving a mean score of 4.3/5, which confirms the system is both accessible and easy to implement across the network.

### 2.5 Preliminary Explainability Analysis

Beyond predictive accuracy, a preliminary post-hoc analysis was conducted to probe whether the morphological features exploited by MosAICo-Net are consistent with established entomological taxonomy, by applying Gradient-weighted Class Activation Mapping (GradCAM++) (Chattopadhay et al., 2018). Such technique projects the gradient of the class score with respect to the final convolutional feature maps back onto the input image to produce a coarse localization heatmap of class-discriminative regions. Although these results are exploratory and constitute a qualitative complement rather than a quantitative evaluation, the patterns observed are sufficiently consistent to warrant reporting.

As shown in Figure 7, the GradCAM++ heatmaps generated for *Anopheles maculipennis s.l.* concentrate activation predominantly on the wings of the specimen. This is directly consistent with its primary di-chotomous key: the characteristic spotted wing pattern from which the species epithet *maculipennis* (Latin: “spotted wing”) derives. The convergence between network attention and established morphological taxonomy was observed across multiple representative specimens, suggesting that this behavior reflects a stable learned representation rather than an artifact of individual images.

**Figure 7:**
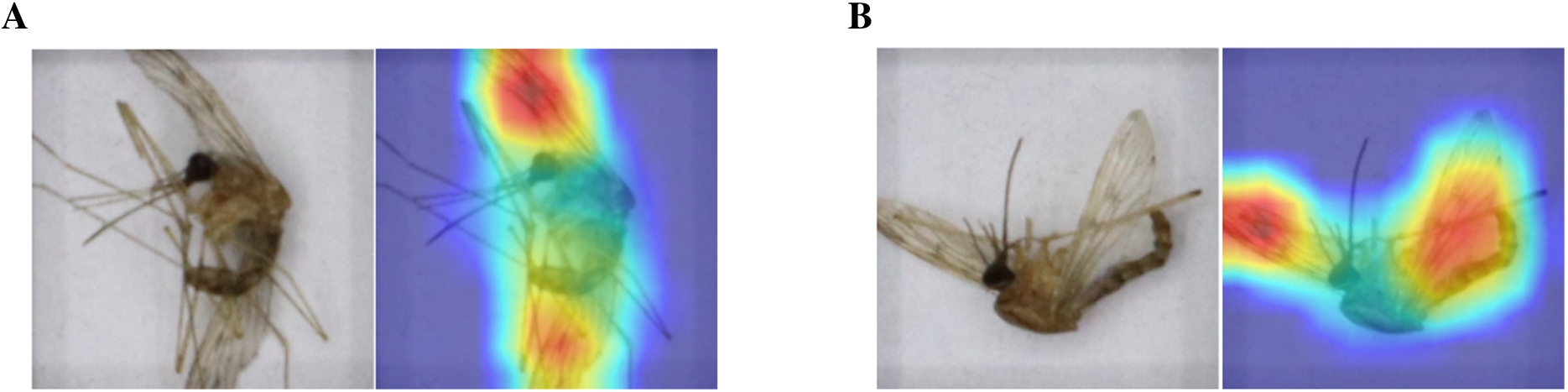
GradCAM++ activation maps for *Anopheles maculipennis s.l.*. Two representative specimen pairs are shown, each displaying the 384 × 384 input image (left) alongside the corresponding GradCAM++ heatmap overlaid on the image (right). In both cases the network’s class-discriminative activations are concentrated on the wings, consistent with the primary dichotomous-key feature used to identify this species. Warmer colours (red/yellow) indicate regions of highest gradient magnitude.

## 3 Material and Methods

### 3.1 The MosAICo Network and Dataset Construction

Data collection was orchestrated through the MosAICo Network, a consortium established to standardize entomological data protocols across Italy, comprising 18 partners across 16 regions, including local health authorities and research institutes.

This collaborative infrastructure enabled a dual-sourcing strategy, leveraging both baseline reference strains from laboratory insectaria, private collections and wild-caught specimens from field operations. By incorporating both pristine, taxonomically verified individuals and damaged, real-world trap catches, the resulting dataset—encompassing 12,499 mosquitoes across 15 distinct species—captures the full spectrum of morphological variance required for robust, species-level discrimination.

While 15 species were initially collected, the training phase was restricted to seven priority taxa, selected based on high sample density and a stringent requirement for each species to represent at least three distinct geographical origins to ensure robust generalization. Notably, *Ae. aegypti* was included as a primary target class due to its critical role as a potential emerging vector in Europe, despite being currently absent from Italy but present in some countries in the Mediterranean basin. This taxonomic selection prioritized both medically relevant vectors and geographically abundant species, ensuring model efficacy against the ‘bycatch’ commonly encountered during routine surveillance activities.

### 3.2 The “Anchor” Strategy for Open-Set Recognition

Conventional deep learning classifiers typically operate under a closed-set assumption, forcing every input into a predefined category. This represents a critical limitation in entomological surveillance, where “bycatch” (non-target species) frequently outnumbers primary vectors. To mitigate this, we implemented a hybrid dataset construction strategy specifically tailored for Modified Evidential Deep Learning (m-EDL) (Nagahama, 2023). Unlike standard approaches that treat “Unknowns” as mere low-confidence predictions, our framework explicitly models background biodiversity as a distinct taxonomic class through a tripartite functional partition (Table S1):

1. **Target Classes (Closed Set)**: The model was trained to identify seven priority vector species: *Ae. albopictus*, *Ae. aegypti*, *Ae. caspius*, *Ae. koreicus*, *Cx. pipiens*, *An. maculipennis s.l.*, and *Ae. vexans*.
2. **Anchor Unknowns (Training Outliers)**: To provide the model with explicit features of irrelevant specimens, a heterogeneous “Unknown” class was curated. This category includes *Ae. geniculatus*, *Ae. mariae*, *Cx. hortensis*, *Ae. detritus*, and *Cx. theileri* (the latter restricted to training due to its single-provenance origin). These “Anchor Unknowns” allow the model to learn the decision boundaries of the background distribution during the training phase.
3. **True Out-of-Distribution (OOD Test Set)**: To rigorously validate the system’s capacity for generalized rejection, specific non-target species—*An. algeriensis* and *Ae. rusticus*—were entirely excluded from the training process. Reserved exclusively for the test set, these specimens serve as a benchmark for True OOD detection, ensuring the model does not merely memorize trained non-targets but successfully rejects taxonomies never encountered during training.

### 3.3 Specimen Localization via Classical Computer Vision

Once acquired a high-resolution image of a mosquito populated cassette, the system proceeds to isolate individual mosquito specimens later passed to the model. Crucially, this localisation stage is not performed by a machine learning detector—typically YOLO (Redmon et al., 2016)—but instead relies on a deterministic, OpenCV-based registration and segmentation pipeline. This design choice reflects the fact that cell positions are entirely determined by the physical grid, whose geometry is fixed and known a priori: exploiting this constraint eliminates the need for a learnable detector and the associated training data burden (Zhong et al., 2018).

The pipeline proceeds in three stages:

1. **Grid registration**: the acquired image is aligned to one of several pre-annotated reference grid templates. Alignment is achieved via Enhanced Correlation Coefficient (ECC) minimization with a full homography motion model (Evangelidis and Psarakis, 2008), estimated on downsampled (600×400) binary masks of the grid lines. The warp matrix producing the highest correlation coefficient (threshold: 0.3) is selected, and the pre-computed bounding boxes of each grid cell are propagated to the acquired image through perspective transformation.
2. **Occupancy detection**: each warped cell ROI is processed independently to determine whether it contains a specimen. After applying a fixed inner margin (62 px) to suppress the grid border, the grayscale inner ROI is binarised using OpenCV’s adaptive mean threshold, followed by morphological erosion and dilation, to consolidate foreground blobs. A connected-component analysis then determines cell occupancy: a cell is flagged as occupied—and its bounding box retained—if and only if at least two connected components are detected (i.e., background plus at least one foreground blob).
3. **Crop extraction**: each occupied cell—whose bounding box may vary in dimensions depending on grid slot size—is extracted as an RGB patch. Because the neural backbone requires a fixed 384×384 pixel input, a random crop of this size is sampled from the patch at inference time, providing a lightweight form of test-time stochasticity consistent with the random-crop augmentation used during training.

### 3.4 The “MosAICo-Net” Architecture

To address the challenges of “Open Set” recognition (Geng et al., 2020), we developed MosAICo-Net, a novel architecture that integrates Modified Evidential Deep Learning (m-EDL) for outlier rejection (Nagahama, 2023) and Conformal Prediction for ambiguity resolution (Angelopoulos and Bates, 2023). Adopting a “Safety First” design philosophy, the pipeline is engineered to prioritize the rejection of unrecognized specimens over the risk of erroneous forced classification.

#### Backbone and Feature Extraction

MosAICo-Net utilizes an EfficientNetV2-M backbone, pre-trained on ImageNet-21K (Tan and Le, 2021), as its primary feature extractor. This configuration generates a 1,280-dimensional representation from the terminal convolutional stage, which is subsequently processed by a single-hidden-layer MLP projection head. The resulting architecture comprises approximately 53 million parameters, maintaining a computational footprint sufficiently lightweight to support CPU-based inference during deployment. To preserve the fine-grained morphological features essential for species-level discrimination, input images are processed as 384 × 384 pixel tiled crops.

#### Outlier Rejection via m-EDL (Red Box)

Conventional softmax classifiers often yield point estimates that mask predictive uncertainty, rendering them inadequate for open-set scenarios. To overcome this limitation, the final classification layer was replaced with a Modified Evidential head (Nagahama, 2023). This module outputs per-class evidence values e, which define the concentration parameters ***α*** = 1 + **e** of a Dirichlet distribution across *K* target species and an additional (*K* + 1)-th “Unknown” category, the latter characterized by a fixed evidence prior *e_u_* = *K*. This formulation enables the model to explicitly decouple aleatoric uncertainty (data-inherent ambiguity) from epistemic uncertainty (model ignorance). A specimen is categorically rejected and assigned to the red box if the (*K* + 1)-th class exhibits the highest predicted probability.

#### Ambiguity Resolution via Conformal Prediction (Yellow Box)

Specimens surpassing the initial rejection stage are processed by an ambiguity resolution module. Here, the “unknown” class probability is nullified, and conformal prediction is applied over the *K* target species to construct a prediction set *C*(**x**). This set is guaranteed to contain the ground-truth label with a marginal probability of at least 1 − *ɛ*, utilizing a calibrated threshold of *τ* = 0.1 and a coverage parameter of *ɛ* = 0.02. The cardinality of *C*(**x**) determines the system output:

1. Green Box (|*C*| = 1): a single species is reported with high confidence.
2. Yellow Box (|*C*| = 2): two candidate species are listed in descending order of probability, flagging ambiguity for human review.
3. Red Box (|*C*| *>* 2): the prediction set is too broad to be informative; the specimen is escalated for expert inspection.

A critical override is implemented for *Ae. aegypti*, currently absent in the Italian context, due to its significant public health implications. Any detection that would otherwise result in a single-species green box for *Ae. aegypti* is automatically downgraded to a yellow box, listing *Ae. albopictus* as the second candidate. This conservative protocol ensures that all putative *Ae. aegypti* identifications undergo mandatory secondary review by an expert.

### 3.5 Training and Validation Protocols

To ensure that the MosAICo-Net evaluation reflects real-world deployment challenges, a rigorous geography-aware partition strategy was implemented. The dataset was partitioned such that, for each species, the held-out test set comprises specimens from geographical provenances entirely disjoint from those in the training set. The sole exception is *Culex theileri*, which was restricted to the training set due to the availability of only a single provenance. This deterministic split prevents “species–location” clusters from straddling the train–test boundary, providing a stringent assessment of the model’s generalization capacity to novel environments. Following this partition, a 5-fold stratified cross-validation was applied to the training corpus. Each fold was further enriched with an auxiliary set of legacy images (characterized by distinct illumination and the absence of an alignment grid) for four target species (*Ae. aegypti*, *Ae. albopictus*, *Ae. koreicus*, and *Cx. pipiens*). This auxiliary corpus facilitates convergence and serves as a form of multi-condition data augmentation, increasing the diversity of photometric contexts encountered during training.

To further bolster robustness against the illumination and positional variability inherent to field-deployed devices, a comprehensive augmentation pipeline was executed using the albumentations library (Buslaev et al., 2020). Transformations were organized into three synergistic domains:

1. Geometric: random grid distortions, horizontal flips (*p* = 0.5), random shifts and rescaling (up to ±20%, with *p* = 0.25), random rotations (up to ±45*^o^*, with *p* = 0.25), affine shears (up to ±20*^o^*, with *p* = 0.25), and random cropping to the 384 × 384 target input size.
2. Radiometric: random brightness and contrast perturbations (brightness ±20%, contrast ±10%, with *p* = 0.25), per-channel RGB shifts (±5 units, with *p* = 0.1), hue–saturation–value jitter (±10 units, with *p* = 0.1), FancyPCA color augmentation, random gamma corrections (*γ* ∈ [40, 100], with *p* = 0.1), and random sharpening (p = 0.10)
3. Noise and blur: additive Gaussian noise (variance *σ*^2^ ∈ [10, 30], with *p* = 0.1) and defocus blur (radius ∈ [0.1, 0.2], with *p* = 0.1).

This multi-family strategy systematically addresses the orthogonal sources of domain shift: geometric variability stemming from specimen orientation and camera positioning, and photometric fluctuations arising from ambient lighting and sensor-specific responses.

### 3.6 The MosAICo Acquisition Device

To achieve rigorous standardization across diverse collection sites, we engineered a custom acquisition unit designed to mitigate environmental variables—such as inconsistent focal distances, non-uniform illumination, and perspective distortion—that typically confound handheld photography (Dasari et al., 2024). The device consists of a benchtop imaging chamber with a rigid chassis that maintains a fixed working distance between the optical sensor and the specimen plane. The imaging system comprises a high-resolution commercial sensor (Canon EOS R100, 24.1 MP) paired with a fixed focal length lens (Canon RF-S 10-18mm, locked at 18mm), calibrated to resolve fine-grained morphological features, including scale patterns and wing venation. To ensure radiometric consistency, the chamber is equipped with a dedicated LED system providing constant “Natural White” (4000 K) illumination, thereby neutralizing the influence of ambient laboratory or field lighting.

Furthermore, the specimen stage is specifically designed to accommodate custom magnetic grids, enabling the simultaneous, non-overlapping disposition of up to 82 individuals.The unit operates as a tethered peripheral controlled via a dedicated workstation interface, allowing for real-time focus verification through a live feed. This interface is fully integrated with the MosAICo cloud platform; upon acquisition, images are uploaded for automated inference and database synchronization. This streamlined workflow ensures that each record is systematically tagged with essential metadata—including geolocation, timestamps, and operator identification—thereby eliminating manual file handling and ensuring data integrity across the network.

### 3.7 Field Validation

In order to validate the in silico metrics against field evidence, ten copies of the device were used at twenty sites in Italy during routine surveillance activities in summer 2025 (Figure 8). This ensured consistent radiometric image capture, regardless of local conditions. A total of 25 high-resolution (6000 × 4000) images were analyzed by the MosAICo system, comprising 1522 collected mosquito specimens. The AI predicted counts were compared against the manual identification performed by expert entomologists.

**Figure 8:**
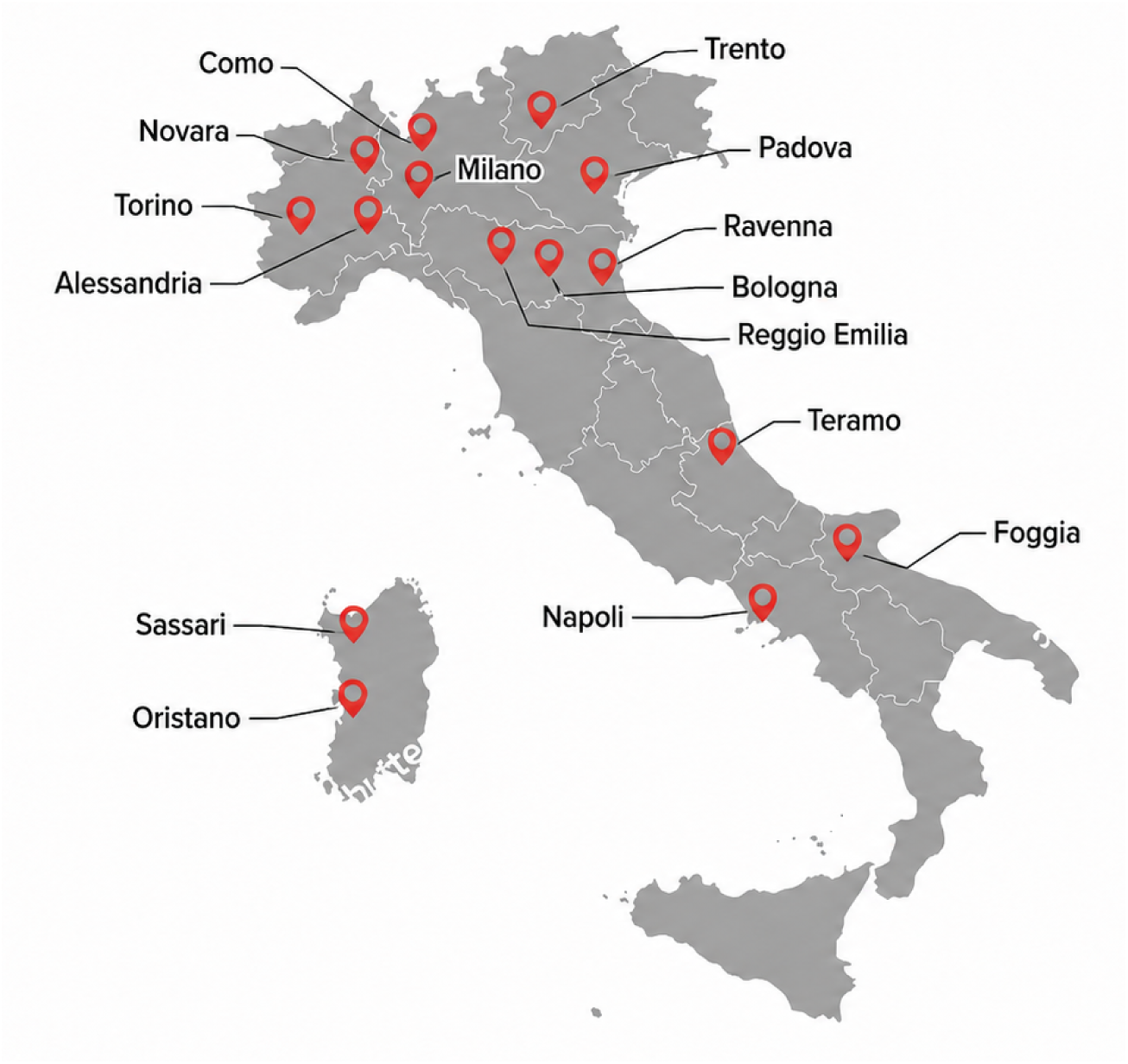
Field collection sites used for real-world validation. Mosquito specimens were collected by collaborating partners from routine field traps distributed across Italy and analyzed with the MosAICo pipeline. The map shows the surveillance sites included in the field-validation dataset, enabling assessment of model performance under operational, real-world collection conditions.

Per-specimen performance was assessed by computing a confusion matrix over the full dataset, from which macro and micro accuracy were derived separately for target species and non-target species. Macro accuracy is defined as the unweighted mean of per-class recall; micro accuracy as the proportion of correctly classified specimens pooled across all classes. Misclassifications were further partitioned into genus-constrained and inter-genus errors.

Population-level agreement was assessed exclusively on green- and yellow-box predictions, as red-box specimens carry no species assignment by design and are routed directly to expert review. For each of the six target species, the automated count (top-1 predictions) was compared against the corresponding manual count. To account for Poisson sampling variability inherent in trap catches, per-species predictive uncertainty was quantified under Jeffreys’ prior as a Negative Binomial, from which 95% prediction intervals and a per-species standard deviation *σ_k_* were derived. Population-level calibration was then assessed via the reduced chi-squared 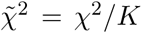, where 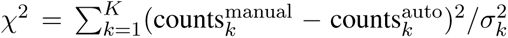 and *K* = 6; values near unity indicate residuals consistent with sampling noise alone.

### 3.8 Cross-Geographic generalization

The geographic generalizability of MosAICo-Net was assessed on 118 *Ae. albopictus* specimens from Ghana, reared under controlled insectary conditions at the University of Pavia. Specimens were organized into four imaging grids, each corresponding to a distinct post-emergence age cohort: 1–2 days (Grid 1, *n* = 35), 7–8 days (Grid 2, *n* = 30), 14–15 days (Grid 3, *n* = 30), and *>*40 days (Grid 4, *n* = 23). All specimens had been identified as *Ae. albopictus* by expert entomologists prior to automated analysis; no model fine-tuning or domain adaptation was performed.

Per-specimen predictions were classified according to the tricolor confidence scheme (Section 3.4), and coverage rates for each category were computed as fractions of the 118 specimens. Accuracy was evaluated at two levels: top-1 accuracy on green-box specimens, and top-2 accuracy on the combined set of green-and yellow-box specimens. To characterize the effect of specimen age on model confidence, coverage and accuracy were stratified by grid, and the species composition of yellow-box prediction pairs was recorded to assess consistency with the confusion patterns documented in the Italian field validation.

## 4 Discussion

MosAICo demonstrates that entomological surveillance can be effectively digitized at a national scale without sacrificing taxonomic reliability. While previous studies have established the theoretical capability of Deep Learning to identify insects (Høye et al., 2021; Teixeira et al., 2023), MosAICo is among the first to successfully operationalize this capability in a distributed, real-world network. By decoupling the acquisition (the device) from the trap, we resolved the image variance issues that have historically prevented comparable systems from achieving the consistency required for reliable operational deployment (Auer-swald and Branscomb, 2003). This validates the hypothesis that hardware standardization is the missing link required to translate in silico accuracy to in situ utility.

A critical distinction between MosAICo and emerging handheld solutions (e.g., VectorCam (Dasari et al., 2024) or VectorBrain (Li et al., 2024)) is our approach to the Domain Shift problem. Deep Learning models degrade rapidly when the inference environment (field lighting, angles) differs from the training environment. Smartphone-based solutions introduce irreducible variance, requiring massive datasets to force generalization. MosAICo takes the inverse approach: by standardizing the acquisition hardware, we physically constrain the inference domain using a rigid hardware standard. This hardware lock-in allows for batch-processing efficiencies that smartphone apps cannot match. While an app scales linearly (one photo per mosquito), MosAICo introduces high-throughput batch dynamics (82 mosquitoes per exposure), increasing data velocity by nearly two orders of magnitude. This throughput is essential for regulatory verification, where “counting mosquitoes” is not merely a census, but a statistically rigorous density estimation required to trigger public health interventions.

Hardware standardisation, however, addresses only the photometric dimension of domain shift. Biological variability—arising from intraspecific morphological variation across geographically distant populations—constitutes a distinct and orthogonal challenge. To probe this dimension, MosAICo-Net was evaluated on a sample of Ghanaian *Ae. albopictus* specimens. Originally native to Southeast Asia, this species has undergone rapid range expansion across sub-Saharan Africa in recent decades, with its establishment in West Africa closely coinciding with multiple dengue fever outbreaks Longbottom et al. (2023). In Ghana specifically, it has demonstrated successful adaptation to local conditions, exhibiting developmental and survival rates comparable to those of the native vector *Ae. aegypti* Akuamoah-Boateng et al. (2026), and its ongoing spread poses a concrete risk of intensified arboviral transmission, including dengue, Zika, and chikungunya Akyea-Bobi et al. (2023). The choice of Ghanaian specimens therefore constitutes a biologically and epidemiologically motivated stress test: a population never encountered during training, situated thousands of kilometres from any training locality, and belonging to an invasive species of pressing public health significance. The strong performance on this out-of-distribution sample suggests that geographically stratified training, a rich augmentation pipeline, and evidence-based uncertainty estimation together confer robustness beyond the photometric domain. This offers a preliminary indication that MosAICo may contribute to automated vector surveillance in regions such as West Africa, where escalating invasive threats and limited expert capacity make scalable, uncertainty-aware identification tools particularly valuable.

The interpretability analysis presented in Section 2.5 provides a further dimension of clinical utility that extends beyond predictive accuracy. First, it constitutes a verification signal that strengthens expert confidence in the model: when the network’s attention aligns with the morphological feature an expert would examine, the prediction becomes an auditable argument rather than an opaque numerical output. This alignment is particularly consequential for acceptance in regulatory and public health contexts, where accountability for misclassification carries non-trivial consequences. Second, and more prospectively, systematic GradCAM analysis across species may reveal discriminative regions not previously codified in formal identification keys. Furthermore, it can indicate that informative signals reside in anatomical structures that taxonomists have historically underweighted. This carries direct pedagogical value, providing a data-driven complement to classical morphological keys and potentially guiding the training of future entomologists by surfacing overlooked discriminative cues. By rendering the model’s internal logic legible to the expert, MosAICo thus functions not only as a classification tool but as an instrument for knowledge discovery—one whose insights can flow bidirectionally between algorithm and human specialist.

While the challenge of validating image-based identification against human experts is well-documented (Høye et al., 2021), we concede that for damaged specimens or cryptic species complexes, human expertise remains irreplaceable. However, the objective of MosAICo is not to replace the entomologists but to scale them via a Human-in-the-Loop framework. Unlike citizen science initiatives (Uelmen Jr et al., 2023; Jain et al., 2024), which often face challenges in data quality control, MosAICo liberates professional bandwidth. While valuable, citizen science datasets often exhibit collection biases and inconsistent sampling effort that can compromise accurate population size estimation if not rigorously modeled (Tran et al., 2021). By focusing expert attention on rare or invasive species, or complementary molecular xenomonitoring (Bigeard et al., 2024).

The simple web interface operationalizes this Human-in-the-Loop paradigm with minimal friction. Entomologists can confirm or correct a prediction simply by clicking on the corresponding cell directly on the image. Beyond individual validation, the platform allows the automatic download of a comprehensive Excel report for any selection of images, containing all the relevant information for an entomologist, that would otherwise need to compile manually from notebooks and spreadsheets. Together, these features dissolve the administrative overhead that has historically acted as a brake on surveillance data throughput, ensuring that expert time is invested in taxonomic judgment rather than data entry.

While MosAICo addresses many surveillance gaps, it is not without limitations. The current hardware unit, while standardized, lacks the extreme portability of smartphone-only solutions, requiring a laptop/workstation and internet connectivity for the complete workflow. Additionally, while the “Closed Set” currently includes 7 priority target species relevant to the Italian context (including *Ae. albopictus* and *Culex pipiens*), expanding this to include other invasive threats requires ongoing data collection. However, this challenge is vastly ameliorated by the system’s integrated feedback mechanism. By allowing field entomologists to validate or correct AI predictions at the single-specimen level via the web dashboard, the system essentially crowdsources “hard” examples. This creates a self-reinforcing active learning loop, where every correction improves the model’s robustness to local variations without necessitating massive, centralized retraining campaigns.

As biological threats accelerate due to global connectivity, the latency inherent in analog surveillance creates a critical vulnerability in health security infrastructures. MosAICo addresses this by integrating entomological data directly into the digital workflow of public health. Rather than merely archiving biodiversity, this system provides the real-time intelligence required to anticipate invasive risks and enable predictive suppression (Abdi et al., 2025; Tsantalidou et al., 2021). By effectively embedding vector control into the proactive management of urban environments, MosAICo offers the cost-effectiveness and temporal resolution necessary to interrupt transmission cycles before they escalate into epidemics (Tozan et al., 2023; Pepin et al., 2013).

## Acknowledgments

The authors gratefully acknowledge Francesco Tucci, Antonio Pazienti, and Luciano Toma for their valuable support and contributions to the development and implementation of the MosAICo ecosystem.

The MosAICo Project is funded by the European Union - NextGenerationEU; Project code PE00000007,Project title “One Health Basic and Translational Actions Addressing Unmet Needs on Emerging Infectious Diseases (INF-ACT). The activity has been funded by the European Union – Next Generation EU under the Italian Ministry of University and Research (MUR) project ECS00000024 “Ecosistemi dell’Innovazione” – Rome Technopole, public call n. 3277, PNRR – Mission 4, Component 2, Investment 1.5.

## MosAICo Working Group Members

The MosAICo Working Group consists of the following members:

***Istituto Zooprofilattico Sperimentale della Puglia e della Basilicata:*** Maria Assunta Cafiero, Maria Grazia Cariglia.

***Fondazione Edmund Mach:*** Annapaola Rizzoli, Daniele Arnoldi.

***Istituto Zooprofilattico Sperimentale delle Venezie:*** Fabrizio Montarsi, Francesco Gradoni.

***Istituto Zooprofilattico Sperimentale della Sardegna:*** Daniele Dedola, Cipriano Foxi.

***Istituto Zooprofilattico Sperimentale della Lombardia ed Emilia Romagna:*** Francesco Defilippo, Annalisa Grisendi.

***Istituto Zooprofilattico Sperimentale Lazio e Toscana:*** Federico Romiti, Adele Magliano.

***Università di Perugia:*** Roberta Spaccapelo.

***Università degli Studi di Napoli Federico II:*** Marco Salvemini, Marianna Varone, Paola Di Lillo.

***C.A.A. (Centro Agricoltura Ambiente) “Giorgio Nicoli” S.r.l.:*** Arianna Puggioli, Alessandro Albieri, Mario Marinozzi.

***Istituto Zooprofilattico Sperimentale dell’Abruzzo e del Molise:*** Matteo De Ascentis, Silvio Gerardo d’Alessio.

***Università di Camerino:*** Guido Favia, Monica Falcinelli.

***ENEA (Agenzia nazionale per le nuove tecnologie, l’energia e lo sviluppo economico sostenibile):*** Riccardo Moretti.

***Istituto Zooprofilattico Sperimentale Umbria e Marche:*** Elisa Antognini, Stefano Gavaudan.

***Università di Milano:*** Sara Epis, Paolo Gabrieli, Marta Villa.

***IPLA S.p.A. – Istituto per le Piante da Legno e l’Ambiente:*** Andrea Mosca.

***Entostudio:*** Andrea Drago, Patrizia Visentin.

## Supplementary Material

### 4.1 Dataset

**Table S1:**
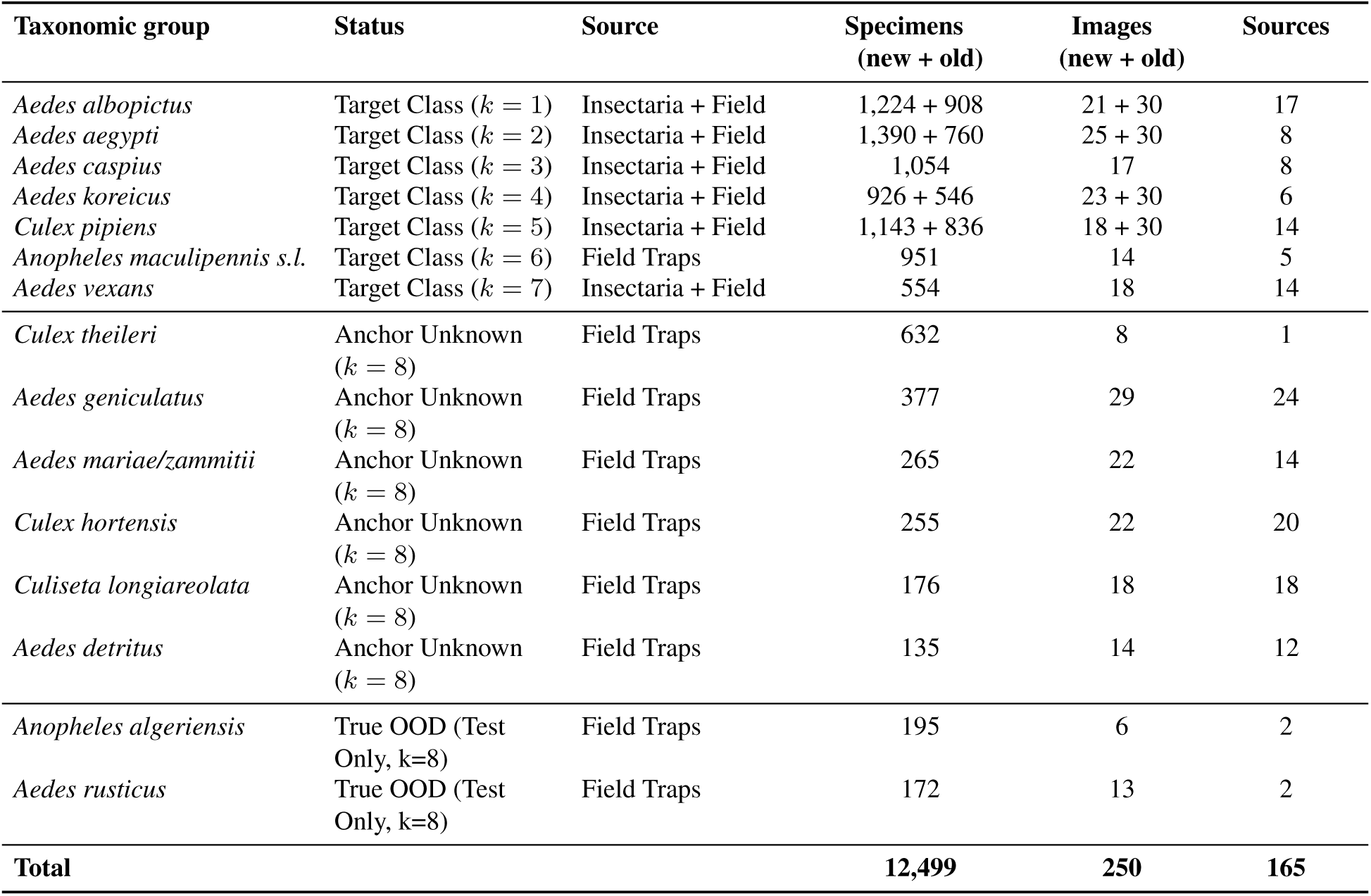
Composition of the MosAICo training and test datasets. Here, *k* denotes the class number assigned to each taxonomic category.

### 4.2 Database

The MosAICo backend is implemented as a relational MariaDB database comprising 15 tables, organized into four functional layers: reference/lookup tables, surveillance metadata, image hierarchy, and AI output and feedback. Figure S1 illustrates the entity–relationship diagram; the tables and their key fields are described below.

#### Reference / Lookup Tables

Seven lightweight tables store controlled vocabularies shared across the schema: `Partners` (institution name), `SamplingTypes` (trap or collection method), `SampleTypes` (specimen origin; e.g. field, insectary), `Channels` (intended data use; e.g. *training* vs. *surveillance*), `Modalities` (image modality type), `Genres` (mosquito genus), and `Species` (mosquito species, linked to `Genres` via `genre id`, with indexes on both `name` and `genre id`). Each table enforces uniqueness on the `name` field.

#### User Management

`Operators` is a one-to-one extension of Django’s built-in `User` model, adding a foreign key to terms-of-service acceptance (`accepted terms`, `accepted terms date`, `accepted terms version`) and `Partners` fields. A database-level `CheckConstraint` enforces that the acceptance date and version must be populated whenever `accepted terms = True`.

#### Surveillance Metadata — Collections

A `Collection` represents one physical trap catch submitted for analysis. Its principal fields are reported in Table S2. Three `CheckConstraints` enforce coordinate validity and that `sampling date` does not postdate the upload `date`.

**Table S2:** Key fields of the Collections table.

| Field | Type | Description |
| --- | --- | --- |
| `camera_name` | `CharField` | Identifier of the acquisition device |
| `date` | `DateTimeField` | Upload timestamp (auto-set) |
| `sampling_date` | `DateTimeField` | Date of field collection |
| `trap_name` | `CharField` | Trap identifier (optional) |
| `sampling_latitude_decimal` | `DecimalField (9,6)` | GPS latitude (-90 to 90) |
| `sampling_longitude_decimal` | `DecimalField (9,6)` | GPS longitude (-180 to 180) |
| `operator_id` | `FK → Operators` | Submitting operator |
| `sampling_type_id` | `FK → SamplingTypes` | Trapping method |
| `sample_type_id` | `FK → SampleTypes` | Specimen origin |
| `channel_id` | `FK → Channels` | Data-use channel |
| `trash` | `BooleanField` | Soft-delete flag |
| `notes` | `CharField` | Optional free-text annotations |
| `app_version` | `CharField` | Client app version at upload |

#### Image Hierarchy — Scenes, Images, SubScenes

Each `Collection` contains one or more **`Scenes`** (individual bulk photographs), each linked to a `collection id`. A `Scene` may be flagged as `flipped` or `trash`, or annotated with `notes`. Each `Scene` has one **`Images`** record, which stores the high-resolution JPEG (`image name`, via `ImageField`) and the raw CR3 camera file (`file name`, via `FileField`), both organised into numbered subfolders to limit filesystem directory size. Uniqueness of filenames is enforced at both the database and application level via a custom `clean()` method. Three computed properties (`secure image url`, `secure file url`, `secure processed url`) resolve storage URLs dynamically without occupying database space.

Each `Scene` is decomposed into one or more **`SubScenes`**, each representing a single grid cell found to contain a specimen by the OpenCV detection pipeline. A `SubScene` stores the bounding-box coordinates of the cell within the parent scene (`bounding box x`, `bounding box y`, `bounding box width`, `bounding box height`).

**Fig. S1:**
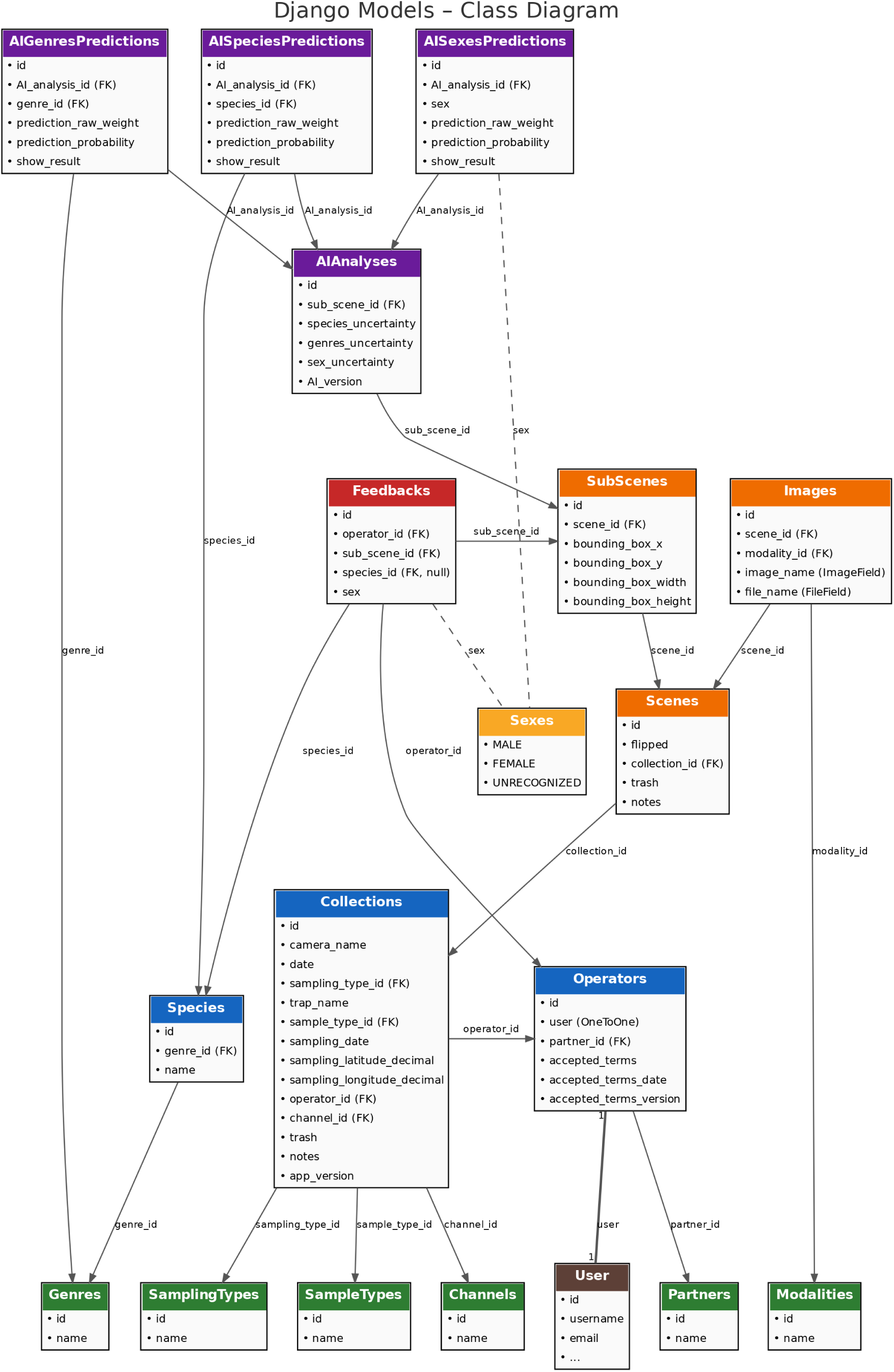
Django class diagram of the MosAICo relational database. Each box represents a database table, with fields listed as bullet points and foreign-key relationships indicated by labelled arrows. Tables are colour-coded by functional layer: Django built-in (brown), Lookup (green), Core (blue), Media (orange), AI pipeline (purple), Feedback (red), Enum (yellow).

#### AI Output — AIAnalyses and Prediction Tables

Each SubScene is linked to one **AIAnalyses** record, which stores aggregate uncertainty metrics (species uncertainty, genres uncertainty, sex uncertainty) and the model version string (AI version).

The predictions themselves are stored in three separate tables, each referencing the parent AIAnalyses via AI analysis id.

- **`AISpeciesPredictions`**: links to `Species` (`species id`), stores the raw m-EDL evidence weight (`prediction raw weight`), the Dirichlet-derived probability (`prediction probability`), and a boolean flag (`show result`) controlling dashboard visibility.
- **`AIGenresPredictions`**: analogous structure at genus level, linking to `Genres` via `genre id`.
- **`AISexesPredictions`**: stores the sex classification (`sex` ∈ {*Male*, *Female*, *Unrecognized*}) with the corresponding `prediction raw weight`, `prediction probability`, and `show result`.

#### Human Feedback — Feedbacks

The **`Feedbacks`** table records expert corrections or confirmations submitted via the web dashboard. Each record links an `operator id` (→ `Operators`) and a `sub scene id` (→ `SubScenes`) to an optional corrected `species id` and/or `sex`. Two database-level constraints ensure data integrity: at least one of `species id` or `sex` must be non-null (`CheckConstraint`), and each operator may submit at most one feedback per `SubScene` (`UniqueConstraint` on `[sub scene id, operator id]`). This table directly feeds the active-learning correction loop described in the Discussion.

